# Central role for fast nociceptors in mechanical nocifensive behavior and sensitization

**DOI:** 10.1101/2025.11.12.688120

**Authors:** John Chwen-Yu Chen, Olivia Le Moëne, Felipe Meira de-Faria, Aikeremu Ahemaiti, Oumie Thorell, Lech Kaczmarczyk, Katharina Henriksson, Jonathan Cole, Håkan Olausson, Walker S Jackson, Saad S Nagi, Malin C Lagerström, Marcin Szczot, Max Larsson

## Abstract

Nociceptors, primary afferent nerve fibers that signal noxious stimuli, are broadly divided into slowly conducting unmyelinated C fibers and fast-conducting myelinated A fibers. Whereas C-nociceptors have been extensively studied, considerably less is known about the function of A-nociceptors. To address this gap, we developed an intersectional genetic approach for robust and selective interrogation and manipulation of mechanically responsive A-nociceptors (A-MNs) in mice. Optogenetic A-MN stimulation induced rapid and precise withdrawal reflexes as well as place aversion and facial expression changes consistent with pain affect, while inhibition strongly impaired mechanical nociceptive withdrawal reflexes, demonstrating that A-MNs are necessary and sufficient for rapid avoidance of noxious mechanical stimuli. Prolonged A-MN activation induced mechanical allodynia and central sensitization. In a rare individual lacking thickly myelinated Aβ fibers, mechanical withdrawal reflexes were completely absent, and pain perception reduced. Together, these findings identify fast-conducting mechano-nociceptors as essential drivers of nocifensive behaviors in mice and humans.

## Introduction

Nociceptors are primary sensory neurons specialized to detect stimuli that damage, or threaten to damage, tissue integrity. Mammalian nociceptors were initially generally considered to be slowly conducting, unmyelinated C fibers, until Burgess and Perl’s unequivocal observation of faster-conducting, myelinated A fiber nociceptors (A-nociceptors) responsive to high-intensity mechanical skin stimulation in the cat^1^. Since then, electrophysiological studies have identified both mechano-selective and mechano-heat responsive populations of A-nociceptor in several mammalian species^2–5^, while single axon reconstructions have provided insight into the anatomy of A-nociceptor terminations in the spinal cord^6–8^. More recently, transcriptomic and other studies utilizing genetic techniques have identified gene markers delineating certain A-nociceptor populations^9–14^.

However, the roles of A-nociceptors in pain and nocifensive behavior, and how they provide input to central sensory pathways, remain poorly understood. For instance, while different populations of C-nociceptors are implicated in nociceptive withdrawal reflexes (NWRs) as well as in affective pain and coping behavior^15–18^, A-nociceptor contribution to these processes is less studied^12, 14^. This can partly be attributed to a dearth of tools for efficient functional manipulation and interrogation of A-nociceptors in animal models. In this study we sought to address this issue by developing a novel intersectional route for genetic targeting of A-nociceptors as a broad population, taking advantage of the unique co-expression of the neurofilament heavy chain (NFH) and the voltage-gated Na^+^ channel Na_V_1.8 in myelinated nociceptors in the mouse^13, 19–22^. We show that this route allows for highly efficient and selective targeting of mechanically activated A-nociceptors, and confirm that such fibers conduct at Aδ to Aβ velocities. We find that they are fundamental for rapid mechanical NWRs but also play a substantial role in affective pain. This is further supported by observations in a rare human Aβ-deafferented individual who failed to exhibit mechanically evoked NWRs and experienced reduced affective pain. Remarkably, prolonged optogenetic stimulation of murine A-nociceptors induced mechanical allodynia as well as both peripheral and central sensitization, revealing a novel route by which pain hypersensitivity can be induced.

## Results

### NFH^CreERT2^;Na_V_1.8^FlpO^ mice enable highly selective targeting of A fiber mechano-nociceptors (A-MNs)

To target a broad population of A-nociceptors in a specific and selective manner, we took advantage of two mouse lines recently developed by us^23, 24^: an NFH^CreERT2^ mouse line expressing tamoxifen-dependent Cre recombinase in cells expressing neurofilament heavy chain (NFH), which in the dorsal root ganglion (DRG) is selectively expressed in myelinated neurons; and a Na_V_1.8^FlpO^ mouse line, which selectively targets FlpO recombinase to neurons expressing Na_V_1.8^FlpO^ (encoded by the *Scn10a* gene), a voltage-gated Na^+^ channel selectively expressed in nociceptors and low-threshold C fiber mechanoreceptors. We reasoned that crossing these mouse lines would direct Cre/Flp-dependent reporter expression specifically to DRG neurons expressing both NFH and Na_V_1.8 (Fig. 1a), and that this targeted group of neurons largely would comprise A-nociceptors. A triple cross of these mouse lines with a mouse line expressing, in a Cre/Flp-dependent manner, the red-activated channelrhodopsin ReaChR^25^ fused to the yellow fluorescent protein mCitrine resulted in membrane-targeted mCitrine expression in numerous DRG neurons. In L3-5 DRGs of these mice (here termed NFH;Na_V_1.8;ReaChR), 94.5 ± 1.6 % (mean ± S.D.; *n*=4 mice) of neurons exhibiting both NFH immunoreactivity and *Scn10a* mRNA were mCitrine^+^; 90.5 ± 3.6 % showed NFH immunoreactivity and 92.5 ± 3.6 % exhibited *Scn10a* mRNA signal, while 86.2 ± 2.8 % of mCitrine^+^ cells were both NFH^+^ and *Scn10a*^+^ (Fig. 1b, c). Thus, this intersectional strategy resulted in targeting of NFH^+^/Na_V_1.8^+^ neurons with high specificity and selectivity. mCitrine^+^ DRG neurons were medium-to-large sized, markedly different from the size distribution of neurons that were *Scn10a^+^* only, but similar to that of NFH^+^ only neurons (Fig. 1d). Notably, only 48.0 ± 2.2 % (*n*=4 mice) of mCitrine^+^ neurons in L3-5 DRGs showed detectable immunoreactivity for calcitonin gene-related peptide (CGRP), a marker of peptidergic nociceptors; conversely, 36.4 ± 4.8 % of CGRP^+^ neurons were mCitrine^+^ (Fig. 1e, f). Few mCitrine^+^ neurons exhibited TRPV1 immunoreactivity (6.0 ± 3.1 %), while essentially no co-localization was observed with binding sites for isolectin B_4_ (IB_4_), a marker of non-peptidergic C fibers.

**Figure 1.**
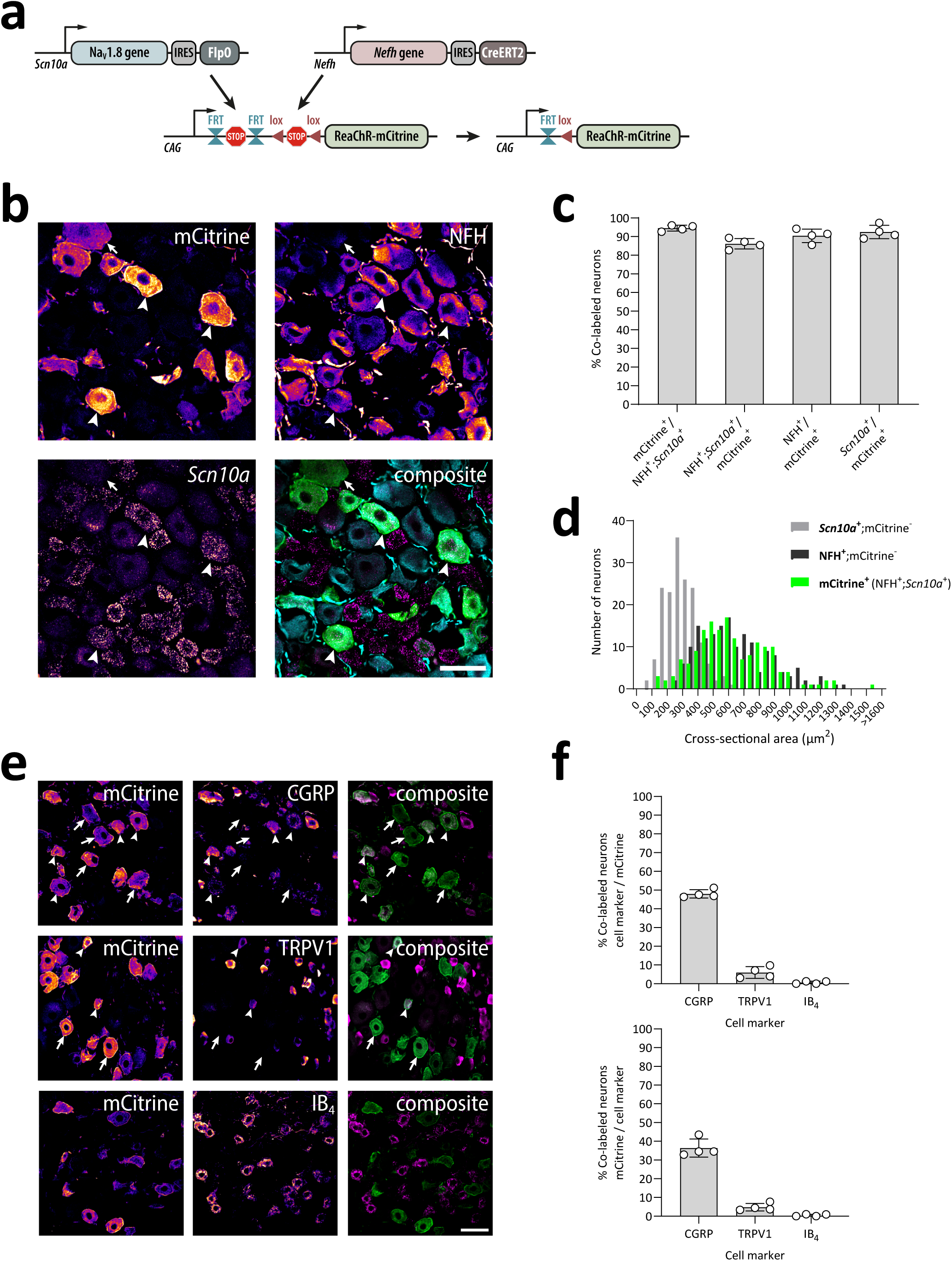
Verification of the NFH;Na_V_1.8;ReaChR mouse line. **(a)** Schematic of the intersectional genetic targeting approach. **(b)** Co-localization of mCitrine immunoreactivity (IR) with *Scn10a* transcript detected using in situ hybridization and NFH immunofluorescence in a lumbar DRG. Single optical section, 20x/0.8 objective. mCitrine, *Scn10a*, and NFH panels are pseudo-colored using the Fire look-up table in Fiji to enhance visibility. Examples of co-localized cells are indicated by arrowheads, while arrows in inset show examples of mCitrine-IR cells with low levels of *Scn10a* transcript. Scale bar, 50 µm. **(c)** Recombination efficiency with respect to *Scn10a*^+^ cells (mean ± SD) in L3-L5 DRGs. Each data point indicates DRG sections from a single mouse; *n* = 4 mice. **(d)** Size histogram of mCitrine^+^, *Scn10a*^+^;mCitrine^−^, and NFH^+^;mCitrine^−^ DRG neurons. *n* = 4 mice (3 ganglia per mouse). **(e)** Co-localization of mCitrine-IR with CGRP-IR, IB4 binding, and TRPV1-IR in lumbar DRGs. Single optical section, 20x/0.8 objective. Scale bar, 50 µm, valid for all panels. **(f)** Quantification of the co-localization of mCitrine with the indicated cell markers. Error bars indicate SD. *n* = 4 mice (3 ganglia per mouse).

Given the medium-to-large size of mCitrine^+^ neurons in the NFH;Na_V_1.8;ReaChR mice and their expected identity as myelinated nociceptors, we decided to examine the size of the peripheral fibers arising from the targeted neurons in transverse sciatic nerve sections. As some but not all of the neurons were found to express CGRP, we also co-immunolabeled the sections for this neuropeptide. Two populations of CGRP^+^ axons were found; one that was associated with mCitrine fluorescence, and one that was not. Axons of the mCitrine^-^/CGRP^+^ population were small, most of them with a diameter less than 0.5 µm, whereas axons of the mCitrine^+^/CGRP^+^ population generally were larger than 1 µm (Fig. 2a). Notably, a population of mCitrine^+^ axons lacked detectable CGRP immunoreactivity, and while the size distribution of this population partially overlapped with CGRP^+^/mCitrine^+^ axons, some of the axons lacking CGRP were larger. To determine the conduction velocities (CVs) of targeted fibers in the NFH;Na_V_1.8;ReaChR mice, we employed an *ex vivo* optogenetics/electrophysiology approach, where optogenetically induced compound action potentials (CAP) were recorded in DRG-dorsal root preparations. First, reference measurements were conducted using electrical dorsal root stimulations. Incrementally increasing electrical stimulations (0 – 2 mA, 0.1ms) to the DRG progressively activated Aβ, δ and C fibers. The fastest CAP was considered to be the Aβ component, followed by the δ and C fiber components^26^. The CV of Aβ fibers ranged from 10.3 to 26.3 m/s with an average value of 20.9 ± 2.7 m/s (Fig. 2b). Aδ fibers exhibited CVs between 6.6 and 8.8 m/s, averaging at 7.3 ± 0.6 m/s, and C fibers ranged from 0.42 to 0.68 m/s, with an average value of 0.54 ± 0.04 m/s. Based on our findings and previous reports^3, 27^, fibers with CVs slower than 1.2 m/s were classified as C fibers, those ranging from 1.2 to 10 m/s were categorized as Aδ fibers, and Aβ fibers were defined by CVs greater than 10 m/s. Unlike electrical stimulation, which instantly delivers activation currents to target cells, optogenetic stimulation introduces an activation delay due to the kinetics of ReaChR^25^. To account for this, CV values of ReaChR-activated fibers were corrected by subtracting the average spike delay of ReaChR (2.27 ms, *n*=6) measured via cell-attached current-clamp recordings (Fig. S1). The corrected CV values (*n*=5) ranged from 1.81 to 22.97 m/s, indicating that optogenetic stimulation of ReaChR activated Aβ and Aδ fibers, but not C fibers (Fig. 2b). Among all ReaChR stimulations, two recordings showed activation of both Aβ and Aδ fibers, while in the remaining recordings only Aδ fiber activation was detected.

**Figure 2.**
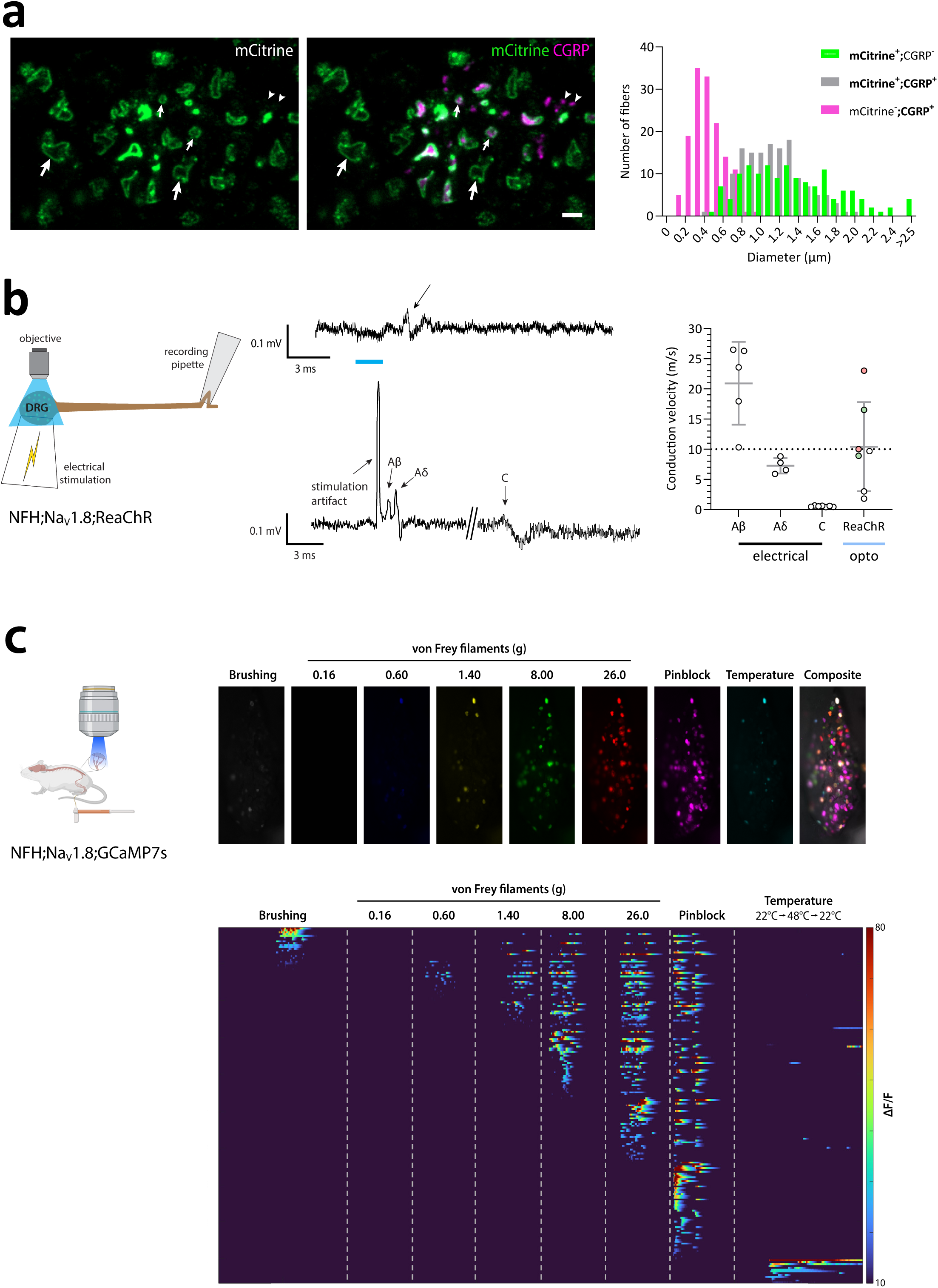
Fiber classes of targeted neurons in NFH;NaV1.8;ReaChR mice. **(a)** Left panels show example micrograph of sciatic nerve sections immunolabelled for mCitrine and CGRP. Single optical section, 63x/1.4 objective. Examples of mCitrine-IR without CGRP-IR axons are indicated by big arrows, axons with colocalization of mCitrine-IR with CGRP-IR are indicated by small arrows, while arrowheads show examples of CGRP-IR axons without mCitrine-IR. Scale bar, 2 µm. Right panel shows size histogram of CGRP^+^;mCitrine^−^, mCitrine^+^;CGRP^-^, and mCitrine^+^;CGRP^+^ axons in sciatic nerve. *n* = 4 mice (2 sciatic nerve sections per mouse). **(b)** Conduction velocity of optogenetically evoked compund action potentials. Left panel shows a schematic of the DRG-dorsal root ex vivo preparation used. Middle panels, example traces of optogenetically (top) or electrically (bottom) evoked recordings. Blue line indicates time of optogenetic activation; arrow in top trace indicates optogenetic response. Right panel, quantification of electrically and optogenetically evoked (ReaChR) CAPs. Green and red color for data points in the ReaChR group indicate that the same-colored points are derived from the same experiments. **(c)** In vivo Ca^2+^ imaging of NFH^+^/Na_V_1.8^+^ DRG neurons. NFH;Na_V_1.8;GCaMP7s mice were subjected to *in vivo* imaging of L4 DRGs to assess responses to natural stimuli applied to the skin of the plantar hind paw. Top panels show example micrographs of responses to different stimuli in an L4 DRG. Bottom panel, a heat map of the responses of GCaMP7s^+^ cells to cutaneous stimuli as indicated. Each row indicates the responses to the applied stimuli of an individual cell. *n* = 193 cells, collected from three L4 DRGs (one ganglion per mouse).

To probe the response characteristics of NFH^+^/Na_V_1.8^+^ fibers to natural stimuli, we performed *in vivo* Ca^2+^ imaging in lumbar (L4) DRGs of NFH;Na_V_1.8;GCaMP7s mice, which expressed GCaMP7s in a Cre/Flp-dependent manner. Very few cells responded to gentle brushing or thermal stimuli applied to the plantar hind paw, whereas the vast majority of cells responded to pin prick applied to the same region of the skin (Fig. 2c). Mechanical thresholds were determined manually using von Frey filaments. Most responsive cells had a moderate-to-high mechanical threshold; no cells were activated by 0.16 g filament, whereas 9 % (17/193) of cells responded to 0.60 g, and 15 % responded to 1.40 g but not 0.60 g filament. About 45 % of cells had a very high threshold, with 20 % responding to 26.0 g but not 8.00 g or lower force filaments, and around 25 % of cells responded only to pin prick. Thus, the population of DRG neurons targeted in these mice largely consisted of A fiber mechano-nociceptors.

### Cutaneous A-MNs form free nerve endings and circumferential hair follicle endings

Next, we sought to determine the skin innervation of mCitrine^+^ fibers. In glabrous hind paw skin, numerous mCitrine^+^ fibers emanated from subepidermal plexa into the epidermis, often extending deep into stratum granulosum (Fig. 3). Some of these fibers formed varicosities along their path through the epidermis, and sometimes, where these could be followed to their distal end, also formed an end bouton. No association with Meissner corpuscle-like structures was found. Some fibers were CGRP^+^, often weakly so, but many were not. In hairy back skin, epidermal CGRP^+^ and CGRP^-^ free nerve endings were found but also circumferential endings around hair follicles. The circumferential endings around hair follicles were CGRP^+^ and likely corresponded to the previously characterized hair pull nociceptors^11, 12^.

**Figure 3.**
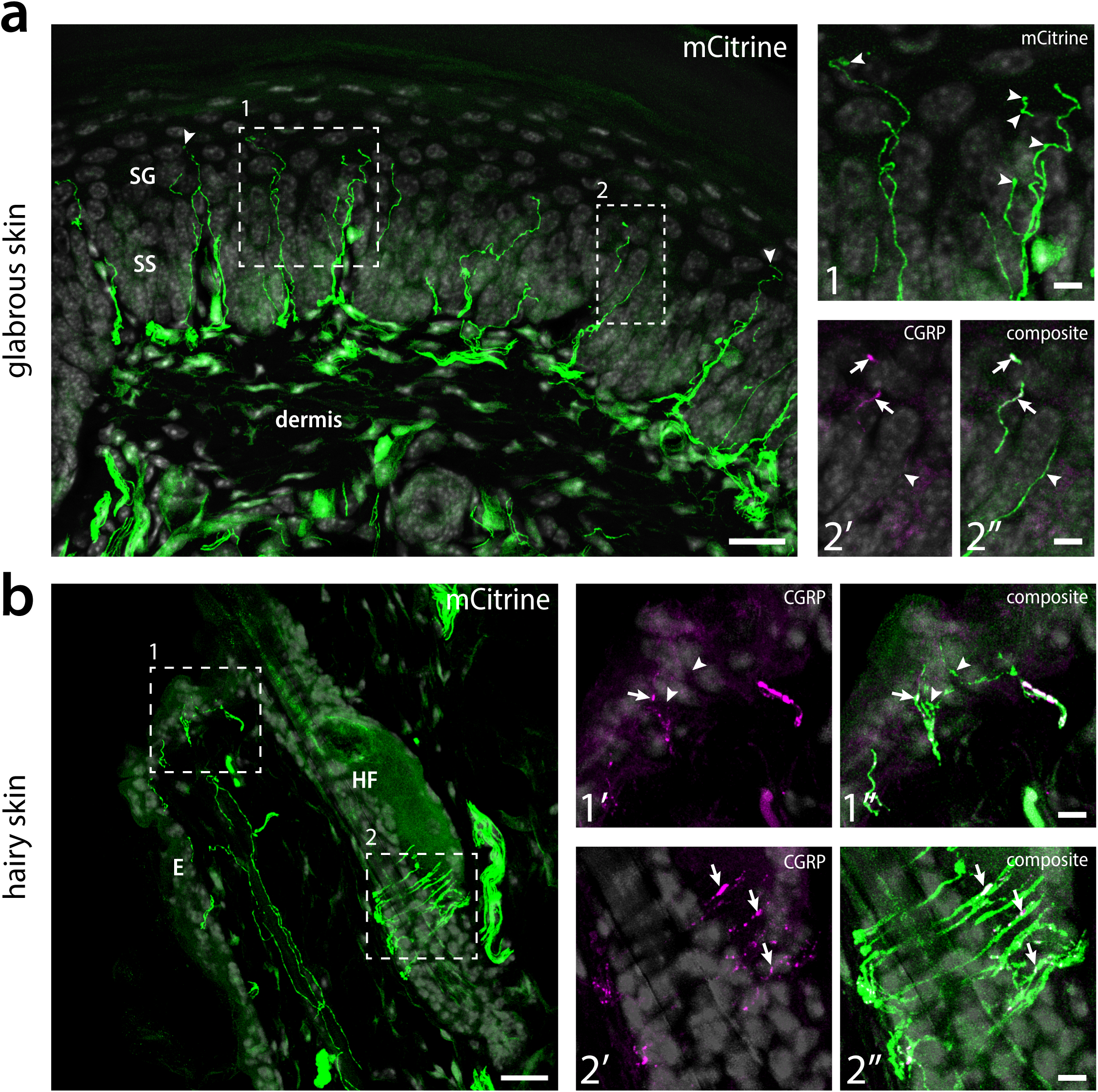
Cutaneous termination of NFH^+^/Na_V_1.8^+^ fibers. **(a)** example micrograph of hind paw glabrous skin from an NFH;Na_V_1.8;ReaChR mouse showing mCitrine^+^ fibers forming free nerve endings in the epidermis. DAPI nuclear staining is shown in grey to visualize layers of the skin. Arrowheads indicate examples of mCitrine^+^ fibers extending through stratum spinosum (SS) deep into stratum granulosum (**SG**). Dashed regions in left panel are shown magnified as numbered in panels to the right. Arrowhead in panel 1 indicate example varicosities in mCitrine^+^ fibers. Panels **2’** and **2’’** show CGRP-IR and mCitrine/CGRP-IR, respectively. Arrows indicate an mCitrine^+^/CGRP^+^ fiber, while arrowhead indicates an mCitrine^+^/CGRP^-^ fiber. Scale bar in left panel, 20 µm; scale bar in right panels, 5 µm. Micrographs are maximum intensity projections of 18 deconvolved optical sections obtained at 0.57 µm separation using a 25x/0.95 objective. **(b)** example micrograph from the hairy back skin of an NFH;Na_V_1.8;ReaChR mouse, showing a circumferential nerve ending around a hair follicle (**HF**) and fibers forming free nerve endings in the adjacent epidermis **(E)**. DAPI staining is shown in grey. Dashed numbered regions are shown magnified in the panels to the right. Panels **1’** and **1’’** show epidermal mCitrine^+^ free nerve endings positive (arrow) or negative (arrowheads) for CGRP. Panels **2’** and **2’’** shows the hair follicle circumferential nerve ending co-localizing with CGRP-IR. Scale bar in left panel is 20 µm; scale bar in right panels, 5 µm. Maximum intensity projection of 20 deconvolved optical sections aqcuired at 0.57 µm separation using a 25x/0.95 objective.

### A-MNs terminate throughout most of the spinal dorsal horn in a segmentally specific manner

In the spinal cord, central processes of targeted mCitrine^+^ neurons were distributed throughout much of the dorsal horn (Fig. 4). The densest innervation was found in lamina I, extending into outer lamina II (II_o_), the ventral border of which was defined by the band of IB_4_ binding. In general, few processes were found in the ventral two-thirds of lamina II; however, in the lumbar enlargement, medial parts of lamina II exhibited considerably denser innervation (Fig. 4b). As the medial dorsal horn in these spinal segments selectively receives input from glabrous skin, this pattern suggests that a glabrous skin-specific myelinated nerve fiber population which uncharacteristically terminates in this region is among those targeted in NFH;Na_V_1.8;ReaChR mice. Furthermore, mCitrine^+^ processes were detected in lamina III and IV albeit at a lower density than in the superficial dorsal horn. A few medial lamina IV fibers appeared to cross the midline to the contralateral dorsal horn, while a few processes entered dorsal lamina X. Somewhat surprisingly however, in lamina V substantial innervation was restricted to the most dorsal part of the lamina, whereas more ventrally only occasional fibers were observed. Moreover, mCitrine^+^ fibers were not uniformly distributed along the mediolateral axis in laminae IV-V, as some of them congregated into large plexa. In sacral spinal cord, in addition to innervation of lamina I and deeper laminae, dense innervation was also observed in the sacral dorsal commissural nucleus (SDC) and in the sacral parasympathetic nucleus (SPN) (Fig. 4c), indicating a large contribution of A-nociceptors to pelvic visceral afferent sensory modalities and to the autonomous regulation of pelvic organs.

**Figure 4.**
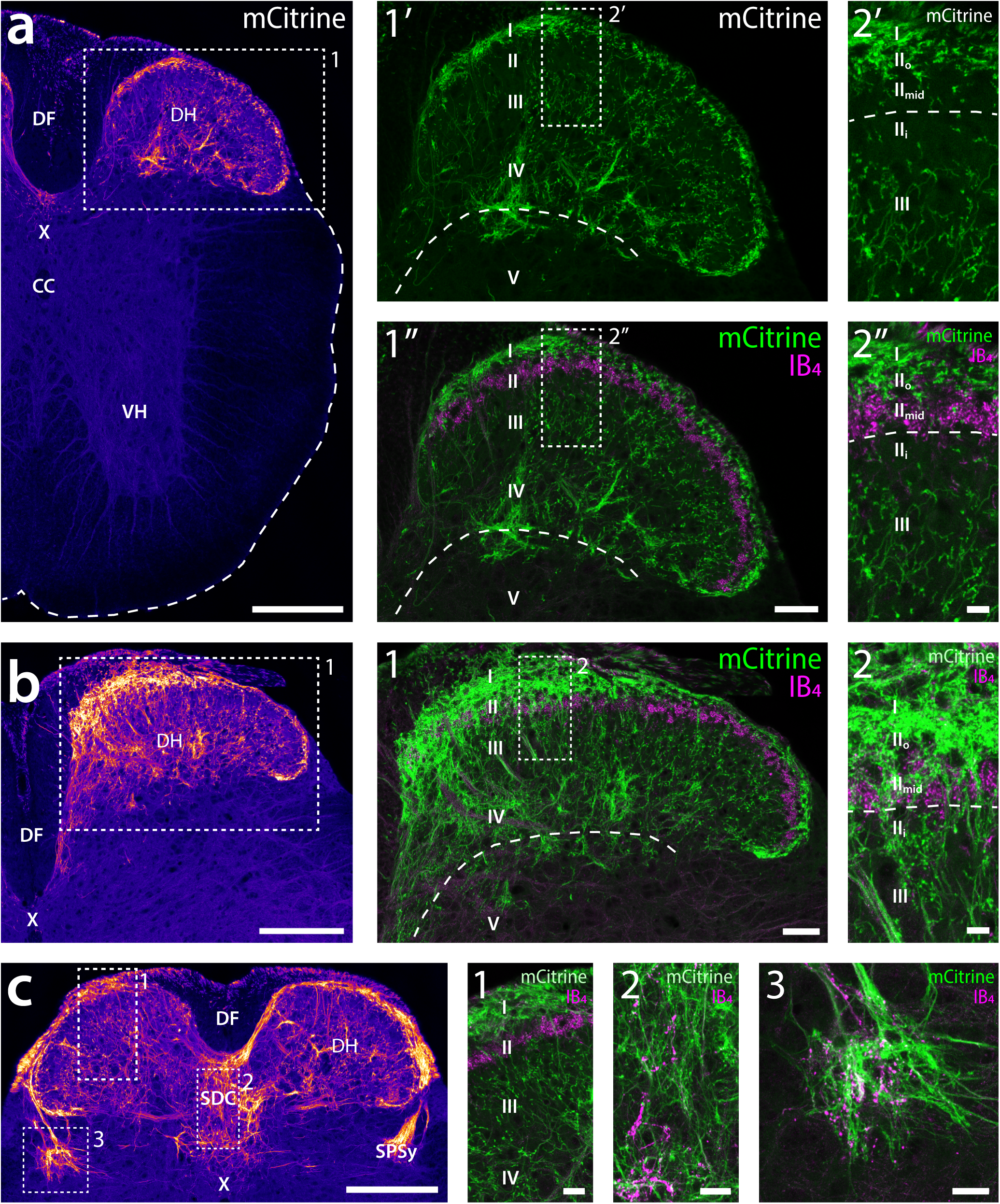
Spinal cord termination of NFH^+^/Na_V_1.8^+^ fibers. **(a-c)** transverse thoracic spinal cord section from an NFH;Na_V_1.8;ReaChR mouse showing endogenous mCitrine fluorescence. The image is pseudo-colored using the Fire look-up table in Fiji to enhance visibility of weak fluorescence. **DF**, dorsal funiculus; **DH**, dorsal horn; **VH**, ventral horn; **CC**, central canal; **X**, lamina X. Note the lack of mCitrine fluorescence in the ventral horn. Dashed region is magnified in the center panels; panel **1’** shows mCitrine fluorescence while panel **1’’** shows mCitrine together with IB_4_ binding (magenta) to outline the middle third of lamina II. Roman numerals indicate Rexed’s laminae. Dashed lines in 1’ and 1’’ show the approximate dorsal border of V. Note the sparsity of mCitrine^+^ fibers in lamina V except the dorsal part, and the comparative density of fibers in lamina IV. Dashed region in 1’ and 1’’ is further magnified in the rightmost panels. Dashed line indicates the ventral border of mid-lamina II (**II_mid_**), as defined by IB_4_ binding. Note the dense plexus of mCitrine^+^ fibers in lamina I and outer lamina II (**II_o_**), the sparseness of fibers in II_mid_ and inner lamina II (**II_i_**), and the intermediate density of fibers in lamina III. Scale bar in left panel, 200 µm; scale bar in center panels, 20 µm; scale bar in right panels, 10 µm. **(b)** transverse section of L4 spinal cord. Note the higher levels of fluorescence in the medial dorsal horn as compared to the thoracic spinal cord in (a). Dashed frame indicates region magnified in center panel (1) which shows both mCitrine fluorescence and IB_4_ binding. Dashed line in the center panel indicates the border between laminae IV and V. Dashed region in the center panel is further magnified in panel 2. Note the much increased density of fibers in lamina II_mid_ and II_i_ as compared to the lateral parts of the same section, as well as to the thoracic spinal cord in (a). Abbrevations and scale bars are as in (a). **(c)** transverse section of S1 spinal cord. Note the dense innervation of the sacral dorsal commissural nucleus (**SDC**) and sacral parasymapthetic nucleus (**SPN**). Dashed frames indicate numbered insets shown in the right panels with endogenous mCitrine fluorescence and IB_4_ labeling. Scale bar in left panel, 200 µm; scale bars in insets, 20 µm. All micrographs are maximum intensity projections of 5 optical sections acquired at 1.04 µm separation with a 20x/0.75 objective.

Primary afferent fibers in the dorsal horn often establish complex synaptic structures known as synaptic glomeruli in the spinal dorsal horn. These glomeruli, in which a central primary afferent terminal makes multiple axodendritic synaptic contacts but also receives inhibitory axo- and dendroaxonic synapses, afford complex signal integration already at the first sensory synapse. However, while these structures are well-established features of C fiber termination^28, 29^, the synaptic organization of A-MNs has only been assessed for a small number of physiologically identified fibers in cat and monkey^30^. Moreover, for these fibers, only terminals in laminae I and V were reported. To determine the synaptic organization of mouse A-MNs and confirm that the presence of mCitrine^+^ fibers in laminae III-IV of NFH;Na_V_1.8;ReaChR mice reflect A-MN synapses in this region, we crossed NFH^CreERT^^2^;Na_V_1.8^FlpO^ mice with an LSL-FSF-APEX2 mouse line, which expresses the peroxidase APEX2 in the mitochondrial matrix in a Cre/Flp-dependent manner. After peroxidase histochemistry, electron dense reaction product is readily detected within mitochondria in APEX2 expressing cells by electron microscopy. In laminae I and II_o_, we observed numerous terminals harboring APEX2^+^ mitochondria; some of these showed a distinct dome-shaped morphology, forming a single axodendritic synapse, while others instead were central terminals of synaptic glomeruli, establishing multiple synapses onto postsynaptic structures (Fig. 5). Notably, these central glomerular terminals were occasionally targets of presynaptic axons forming symmetric, presumed inhibitory^31^, synapses onto the central terminal. Dense core vesicles were found in some terminals in the superficial dorsal horn. In laminae III and IV, APEX2^+^ terminals were also observed, although more sparsely. These terminals were often central terminals of glomeruli, but especially in lamina IV many terminals, despite their generally large size, were found to establish single synapses onto postsynaptic dendrites. As we did not inspect individual APEX2^+^ terminals in serial sections, we could not assess the true number of synapses formed by these terminals.

**Figure 5.**
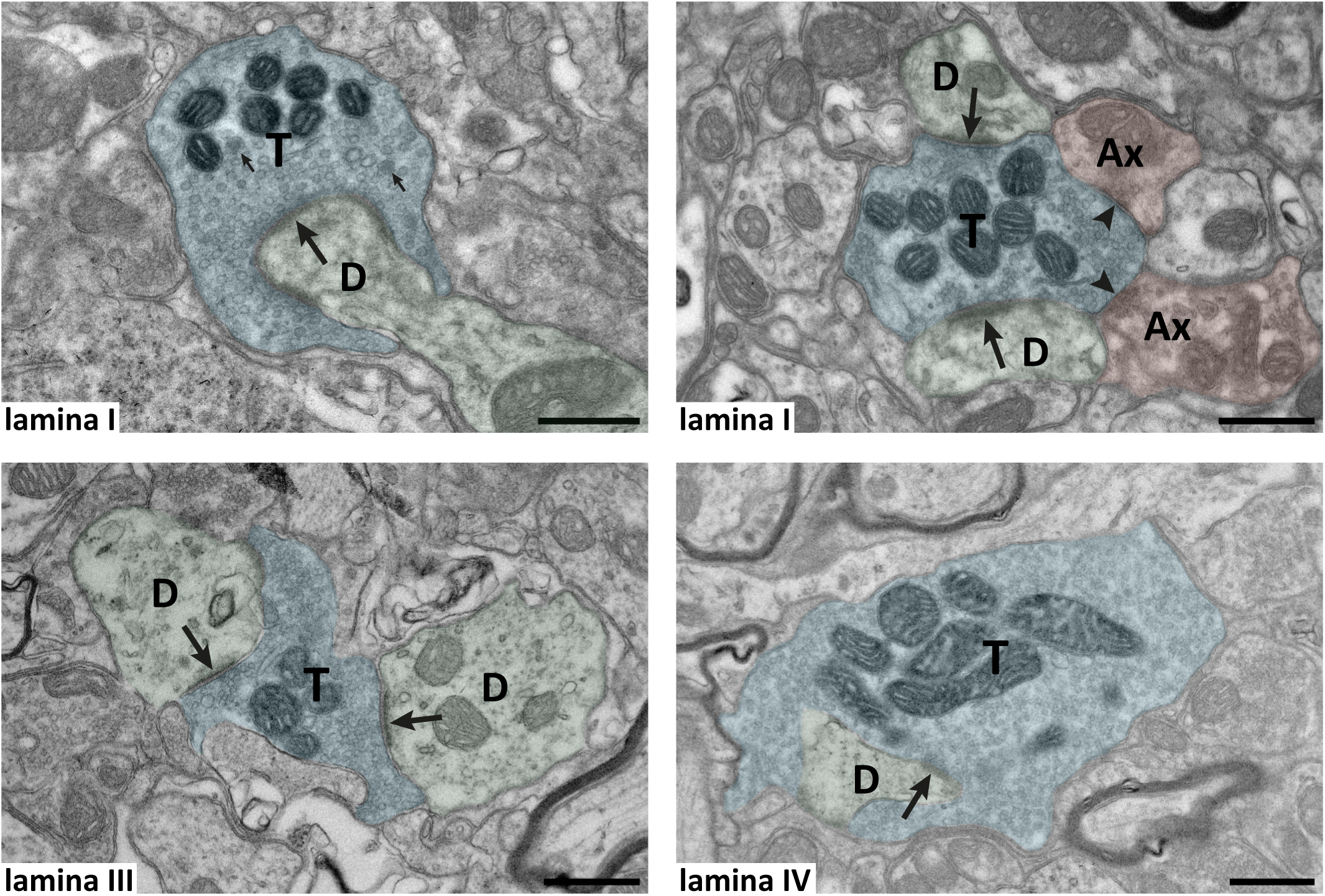
Ultrastructure of spinal termination of NFH^+^/Na_V_1.8^+^ fibers. Shown are example electron micrographs from lumbar spinal cord of NFH;Na_V_18;APEX2 mice. Mitochondria of APEX2 expressing cells show electron dense reaction product in the matrix. In all panels, nerve terminals labeled (**T**) possess mitochondria with APEX2 reaction product that makes those clearly darker in appearance than mitochondria in surrounding neuropil, thus identifying their parent terminal as originating from NFH^+^/Na_V_1.8^+^ fibers. Top left panel shows an APEX2^+^ terminal in lamina I with a simple dome-shaped appearance forming a single synapse (**large arrow**) onto a dendrite (**D**). **Small arrows** indicate examples of dense core vesicles that presumably contain neuropeptides. Top right panel, an APEX2+ terminal in lamina I, that is a central terminal of a synaptic glomerulus, forming two synapses (**arrows**) onto two dendrites but also receiving synapses (**arrowheads**) from two presumed inhibitory presynaptic axons (**Ax**). Bottom left, an APEX2^+^ terminal in lamina III forming a synaptic glomeruli with synapses onto two dendrites. Bottom right, a moderately large APEX2^+^ terminal forming a single obliquely sectioned synapse onto a dendrite. Scale bars in all panels are 500 nm. In all panels terminals and dendrites are pseudo-colored to increase readability.

### A-MNs are required and sufficient for precise nociceptive withdrawal reflexes

Next, we employed NFH;NaV1.8;ReaChR mice to assess behavioral responses to optogenetic stimulation of NFH^+^/Na_V_1.8^+^ fibers. Notably, a single pulse of light applied to the plantar hind paw resulted in a very rapid withdrawal of the paw; using a high-speed camera, we measured the latency to withdrawal from the start of the pulse to 21 ± 3 ms (mean ± S.D.; Fig. 6a). To further characterize the role of these fibers in withdrawal reflexes, we generated NFH;Na_V_1.8;hM4D_i_ mice, where the inhibitory chemogenetic receptor hM4D_i_ is expressed in NFH;Na_V_1.8 fibers. In control mice (positive for NFH^CreERT2^ and hM4D_i_ alleles but negative for Na_V_1.8^FlpO^), i.p. injection of the hM4D_i_ agonist CNO did not affect the mechanical threshold to von Frey filaments (Fig. 6b). However, in mice triple heterozygous for the NFH^CreERT2^, Na_V_1.8^FlpO^ and hM4D_i_ alleles, CNO administration resulted in a marked increase in mechanical threshold. To corroborate this finding, we injected neonatal NFH;Na_V_1.8 mice with an AAV coding for Cre/Flp dependent tetanus toxin light chain (TeTxLC). Compared to NFH;Na_V_1.8 mice injected with a control AAV, these mice when tested as adults had a similarly increased mechanical threshold to von Frey filaments applied to plantar hind paw (Fig. 6c). In contrast to these effects on von Frey thresholds, no effect of chemogenetic inhibition was found with respect to latency to nocifensive responses in the hot plate test (Fig. 6d).

**Figure 6.**
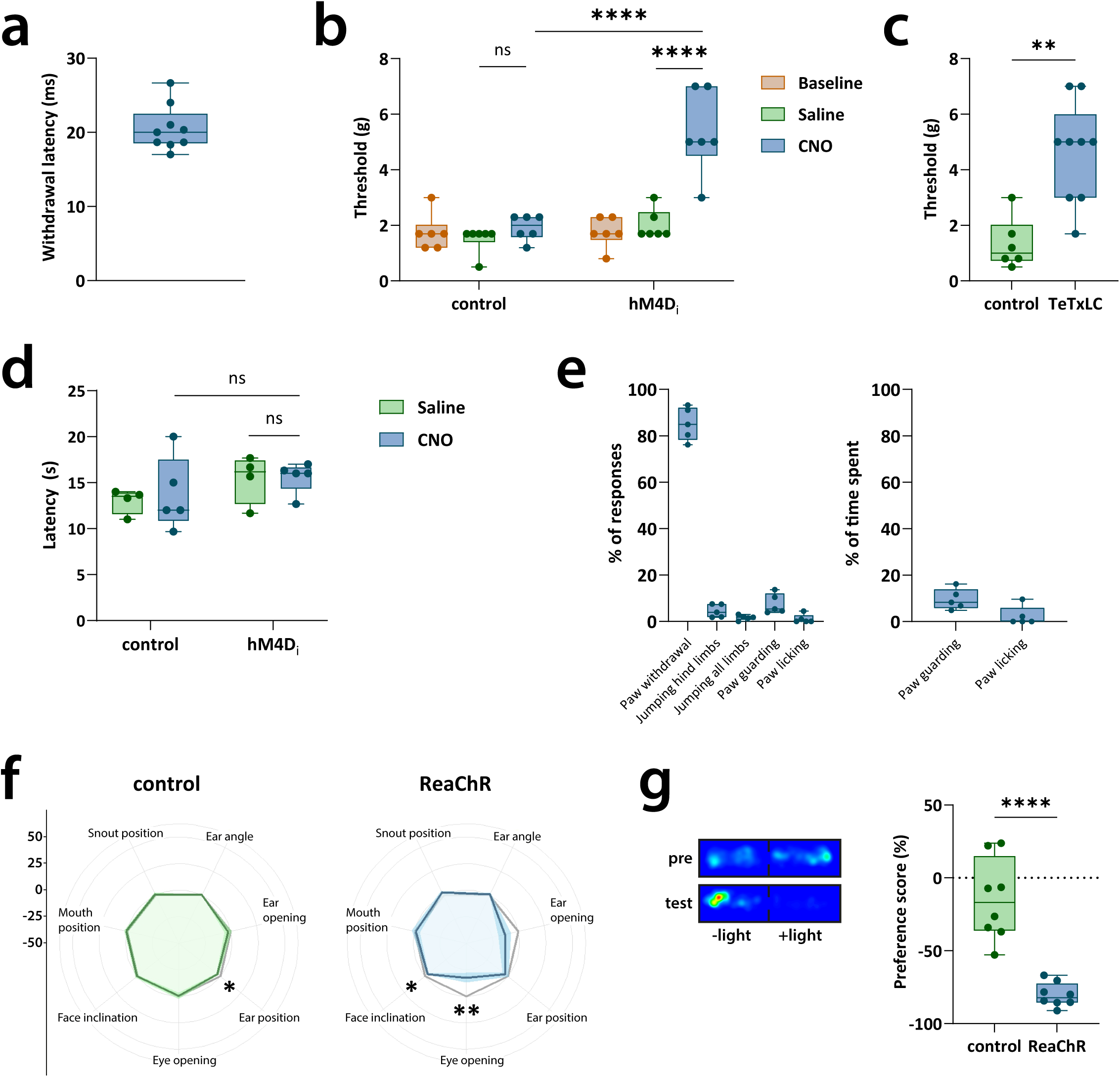
NFH^+^/Na_V_1.8^+^ fibers in nocifensive behavior. **(a)** Paw withdrawal reflex using optogenetic stimulation. For NFH;Na_V_1.8;ReaChR (triple heterozygous) mice, latency to paw withdrawal was 20±3 ms. Control mice did not withdraw their paws (not shown). *n* = 9 NFH;Na_V_1.8;ReaChR mice, *n* = 9 control mice. **(b,c)** Mechanical withdrawal reflex thresholds after inhibition of NFH^+^/NaV1.8^+^ fibers. **(b)** NFH;Na_V_1.8;hM4D_i_ (triple heterozygous) and control mice were injected with either CNO or saline. NFH;Na_V_1.8;hM4D_i_ mice showed significant higher thresholds of paw withdrawal after CNO injection. *n* = 6 NFH;Na_V_1.8;hM4D_i_ mice, n = 6 control mice. ****, *p*<0.0001; ns, p>0.05; two-way ANOVA followed by Tukey’s *post hoc* test. Data passed Shapiro-Wilk normality test. **(c)** neonatal NFH;Na_V_1.8 (double heterozygous) mice were injected with AAV vectors expressing Cre/Flp-dependent TeTxLC or a control plasmid. TeTxLC expressing mice showed significant higher thresholds of paw withdrawal. *n* = 9 TeTxLC mice, *n* = 6 control mice. **, *p*=0.0016; Mann-Whitney two-tailed test. **(d)** Hot plate assay. Each mouse was administered either saline or CNO. No significant difference of latencies to mouse licking, flicking or jumping was shown between NFH;Na_V_1.8;hM4D_i_ and control mice. *n* = 9 NFH;Na_V_1.8;hM4D_i_ mice (4 saline, 5 CNO), *n* = 9 control mice (4 saline, 5 CNO). ns, *p*>0.05, two-way ANOVA followed by Tukey’s *post hoc* test. **(e)** Nocifensive behavior induced by optogenetic stimulation of the plantar hind paw. Left, percentage of responses of each type relative to all observed responses. Right, percentage of time spent guarding or licking the stimulated paw. *n* = 5 mice. **(f)** Facial expression analysis. *n* = 8 NFH;Na_V_1.8;ReaChR mice, *n* = 9 control mice. *, *p*<0.05; **, *p*<0.01; one-sample t-test against baseline. **(g)** Real-time place preference. NFH;Na_V_1.8;ReaChR mice showed less time spent in the light-on chamber. *n* = 8 NFH;Na_V_1.8;ReaChR mice, *n* = 8 control mice. Bottom panel shows an example heat map of time spent in the light-on and light-off chambers for an NFH;Na_V_1.8;ReaChR mouse. ****, *p*<0.0001; unpaired two-tailed Student’s t-test.

It has been suggested that selective activation of myelinated nociceptors may trigger exaggerated nocifensive responses including jumping with both hind limbs or all four limbs, and that normal, well-coordinated nociceptive withdrawal reflexes restricted to the stimulated hind paw require simultaneous activation of A-LTMRs^14^. However, we found that continuous optogenetic hind paw stimulation of NFH^+^/Na_V_1.8^+^ fibers during a period of 1 min yielded primarily simple, ipsi-/unilateral withdrawal reflexes. Of all nocifensive responses, 85 ± 7 % (mean ± S.D., *n* = 5 mice) were such well-coordinated, unilateral reflexes; 5 ± 3 % of responses involved jumping with hind limbs, and 2 ± 1 % of responses comprised jumping with all limbs (Fig. 6e). Further, 8 ± 4 % of responses were guarding of the ipsilateral hind paw whereas 1 ± 2 % of responses consisted of paw licking; only two of five mice exhibited paw licking behavior during the time period scored. Guarding and licking episodes covered 9.6 ± 4.5 % and 2.4 ± 4.2 % of the time, respectively. Thus, we conclude that NFH^+^/Na_V_1.8^+^ fibers are sufficient and necessary for normal, rapid mechanical nociceptive withdrawal reflexes, but not heat nocifensive behavior.

### A-MNs can mediate affective pain

To probe a putative role of NFH^+^/Na_V_1.8 fibers in affective aspects of pain, we first used our recently developed tool for facial expression analysis with optogenetic stimulation of plantar hind paw skin^32^. We detected subtle yet significant changes in facial expression reminiscent of a “pain face” (Fig. 6f). Qualitative inspection of recorded videos showed that salient pain-associated features, such as orbital tightening, were induced by optogenetic stimulation, but occurred only intermittently during the stimulation period. This was likely partly due to the nature of the stimulus; since the manual stimulation of the hind paw resulted in rapid withdrawal and paw guarding/licking, the stimulation was not held at constant intensity or location throughout the stimulation period. To further substantiate a role for these fibers in affective pain, we performed a real time place preference assay (RTPP) in NFH;Na_V_1.8;ReaChR mice. Unlike littermate mice, these mice showed a strong aversion to the light-paired chamber (Fig. 6g), confirming that these fibers were able to activate supraspinal pathways mediating affective pain.

### A-fibers are required for NWRs and normal pain sensation in humans

The above observations indicate that A-MNs are essential for mechanically evoked NWRs. To further test this notion, we turned to a human individual (a 70-year old male) with a selective Aβ deafferentation and sparing of C-fibers^33^. This individual developed a sensory ganglionopathy from C3 level downwards and completely lacks Aβ fibers including Aβ-mechano-nociceptors, which in humans is the only known type of A-MN^34^. We used a custom-made device for rapid punctate mechanical stimulation of the sole of the foot with variable stimulus intensity, and randomized timing to avoid habituation or descending suppression (Fig. 7a). NWR responses were recorded using electromyography (EMG) of the tibialis anterior muscle. Remarkably, mechanical stimulation failed to evoke an NWR at any stimulus intensity up to the individual’s pain tolerance [0-10 visual analog scale (VAS) > 7], unlike in an age-matched control (female, 77 years) (Fig. 7a). Furthermore, mechanical pain threshold (assessed using the same device) was substantially higher in the deafferented subject than in the age-matched control and other healthy control individuals (Fig. 7c); the VAS pain rating at pain threshold was somewhat higher. This is in line with the higher mechanical threshold of C-versus A-nociceptors^34^. Importantly, temperature detection thresholds in the Aβ-deafferented individual were intact, and motor conduction only slightly impaired (Fig. S2). Thus, Aβ nociceptors are critical for mechanical NWRs in humans.

**Figure 7.**
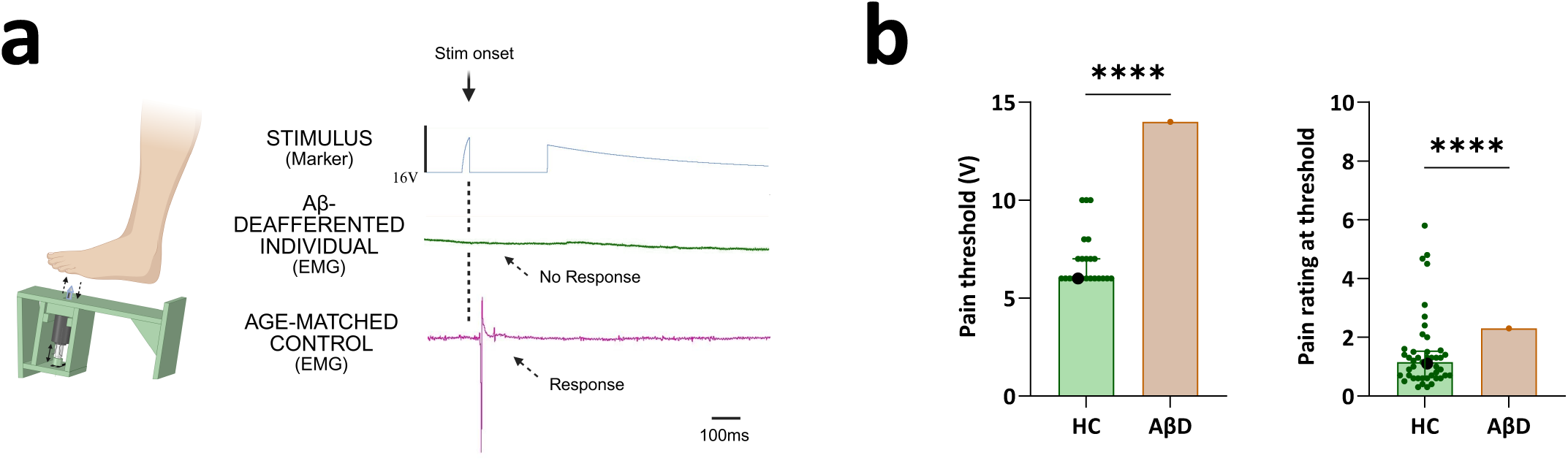
Human nociceptive reflexes and pain psychophysics in healthy control subjects and an Aβ-deafferented individual. **(a)** Mechanically evoked nociceptive withdrawal reflexes were absent in an individual lacking Aβ fibers. *Left*, experimental setup for rapid mechanical stimulation of the sole of the foot. NWR response was assessed using electromyography (EMG) of m. tibialis anterior. *Right*, example EMG traces from the Aβ-deafferented individual and an age-matched control. Top trace shows the stimulus (Stim) represented as voltage applied to the mechanical NWR apparatus. Note that an NWR response is completely absent in the Aβ-deafferented individual. (**b**) Pain threshold (*left*) and ratings (*right*) of rapid mechanical stimulation in the Aβ-deafferented individual (AβD) and healthy control subjects (HC; *n*=23). Large black circles indicates the age-matched control. ****, *p* < 0.0001; one-sample t-test against the Aβ-deafferented individual.

### A-MNs can induce mechanical allodynia and central and peripheral sensitization

Finally, we determined if strong, prolonged selective stimulation of NFH^+^/Na_V_1.8^+^ fibers could induce sensitization. Indeed, after a 5 min long optogenetic stimulation (20 Hz) of plantar hind paw skin (administered during isoflurane anesthesia), the mechanical threshold to von Frey filaments after awakening was substantially reduced compared to the contralateral hind paw, while the response rate to soft brushing of the stimulated paw was increased, indicating that prolonged stimulation of NFH^+^/Na_V_1.8^+^ fibers induced both static and dynamic mechanical allodynia (Fig. 8). Further, the light intensity of an optogenetic stimulus needed to evoke paw withdrawal was reduced, suggesting that the sensitization in part was mediated by increased excitability of NFH^+^/Na_V_1.8^+^ fibers. However, prolonged optogenetic stimulation also induced prominent phosphorylation of extracellular signal-regulated kinase (pERK), a well-established marker of central sensitization, in the superficial dorsal horn of the spinal cord (Fig. 8c). Thus, allodynia induced by prolonged stimulation of NFH^+^/Na_V_1.8^+^ fibers could be mediated by both peripheral and central sensitization.

**Figure 8.**
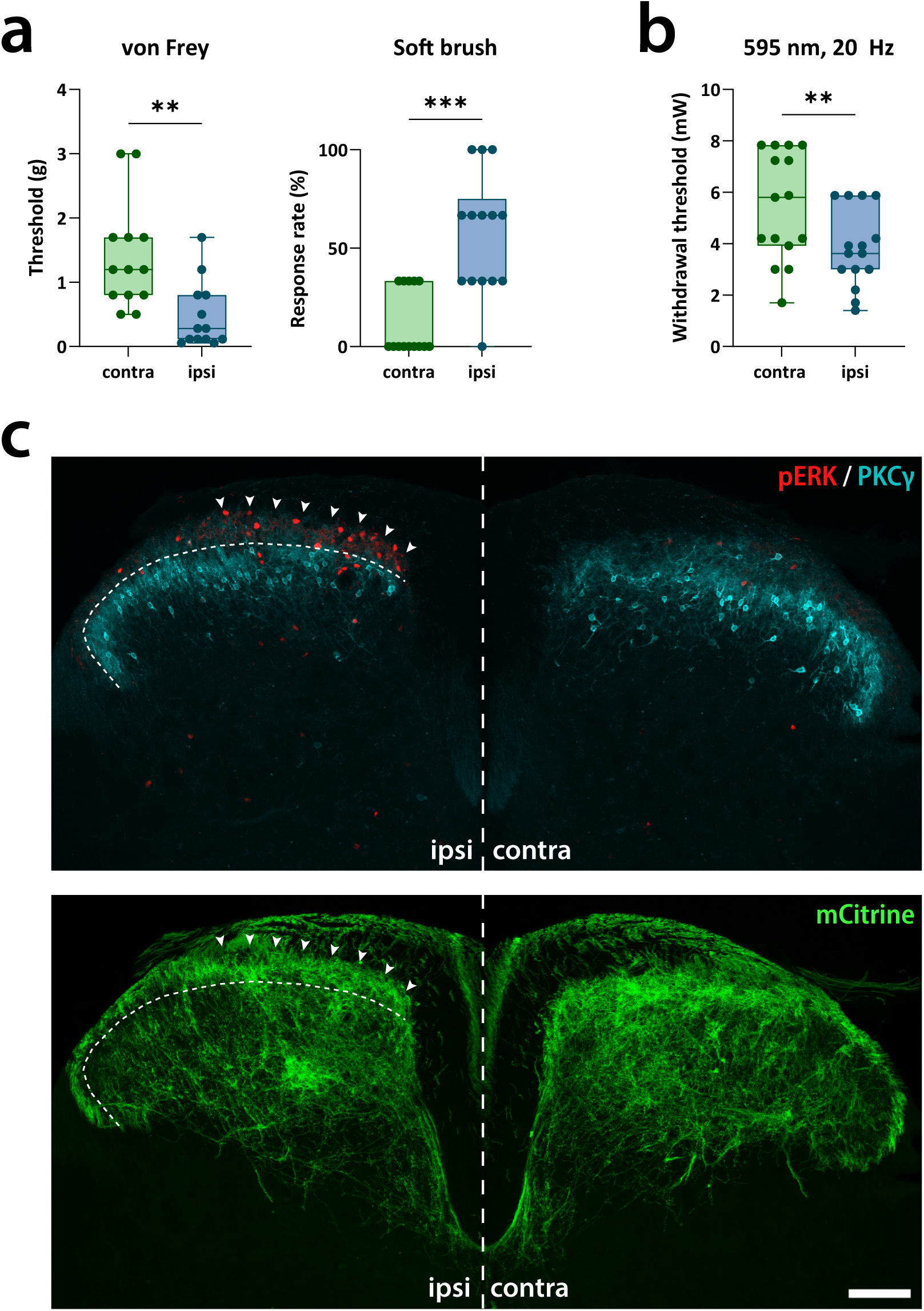
Sensitization by prolonged optogenetic stimulation of NFH^+^/Na_V_1.8^+^ fibers. Optogenetic stimulation (465 nm, 20 Hz, 5 min) was applied to the plantar hind paw of NFH;Na_V_1.8;ReaChR mice. Paw withdrawal reflexes were then assayed with respect to **(a)** mechanical stimulation (*left*, von Frey assay; *right*, soft cotton brush), and (**b**) optogenetic stimulation (595 nm, 20 Hz). Hind paws subjected to prolonged optogenetic stimulation (**ipsi**) were shown to have significantly decreased thresholds compared to the non-stimulated (**contra**) paws. *n* = 13 NFH;Na_V_1.8;ReaChR mice. **(c)** Immunostaining of phospho-ERK (pERK) in the lumbar spinal cord after optogenetic stimulation of the left plantar hind paw. *Top*, pERK and protein kinase Cγ (PKCγ) immunolabeling in a transverse L4 spinal cord section. Note pERK^+^ cells concentrated in a somatotopically appropriate manner in the medial superficial dorsal horn (indicated by arrowheads) receiving input from the hind paw plantar skin. *Bottom*, NFH^+^/Na_V_1.8^+^ fiber innervation in the same section indicated by mCitrine fluorescence. Thin dashed lines indicate border between lamina II_i_ and lamina II_o_ as determined from the dorsal border of the PKCγ immunoreactive band. Vertical dashed lines separates the ipsi-(**ipsi**) and contralateral (**contra**) dorsal horn. Micrographs are maximum intensity projections of 15 optical sections acquired at 1.04 µm separation with a 20x/0.75 objective. Scale bar, 100 µm.

## Discussion

We employed a novel intersectional genetic targeting approach to visualize and manipulate myelinated nociceptors as a broad population in mice. This contrasts with most earlier studies, which have often involved electrophysiological recordings of single fibers, sometimes combined with morphological or neurochemical characterization^1, 6, 7, 22, 35–38^; these studies have by their nature limited sample sizes and were not designed to assess the function of A-nociceptors with respect to pain behavior. More recently, three subpopulations of A-MN were identified in mice using genetic targeting approaches^9, 11, 12, 14^. One of these populations forms hair follicle endings and has been shown to mediate hair pull pain^11, 12^, while another population, targeted using an *Npy2r*^Cre^ mouse line, may be involved in pinprick mediated withdrawal reflexive behavior^14^. A third type of A-MN, characterized by expression of *Smr2*, has recently been described, but its role in nociception and pain behavior is yet unknown^9^. Interestingly, all A-nociceptor populations identified so far express CGRP. In the present study we found that only ∼50 % of NFH^+^/Na_V_1.8^+^ A-nociceptors have this neuropeptide at detectable levels (in line with previous observations in guinea pig^37^). This strongly suggests that one or more populations of A-nociceptor remain unidentified. A-MNs are assumed to primarily form epidermal free nerve endings. Perl and co-workers used the spatial correlation of *post hoc* identified cutaneous nerve endings in cat hairy skin with the receptive fields of electrophysiologically recorded A-MNs to support this notion^38^. However, more direct evidence has been sparse. *Npy2r*^Cre^-targeted CGRP^+^ A-nociceptors were indeed found to form free nerve endings in the epidermis^14^, while another CGRP^+^ population instead formed circumferential nerve endings around hair follicles^12^. Here we confirmed that while certain CGRP^+^ A-MNs form circumferential endings, others that express this peptide form free nerve endings in the epidermis of both hairy and glabrous skin. In addition, we found that A-MNs that lack CGRP also form epidermal free nerve endings in hairy and glabrous skin. No other type of nerve ending was observed in the skin of these mice.

Conventionally, A-MNs have been believed to mainly terminate in lamina I and, to a lesser extent, lamina V^7, 39^. By contrast, Boada and Woodbury found extensive arborization of some A-MNs throughout the dorsal horn^6^. Our present observations suggest that A-MNs provide the most extensive targeting in lamina I and lamina II_o_, while laminae III-IV are more sparsely innervated, and lamina V largely devoid of innervation, except for its dorsal most part. Moreover, the innervation of lamina II appears to be dependent on peripheral input, as only regions of ventral lamina II receiving input from glabrous skin have substantial A-MN innervation. While our observations may appear at odds with those of Boada and Woodbury, re-examination of the reconstructed fibers shown in their study suggest that those fibers that project deeply in the dorsal horn in fact show a rather restricted journey into lamina V, with the most dense arborization in laminae III-IV^6^. Thus, we argue it is warranted to revise the general view of the spinal termination of A-MNs, such that these fibers terminate primarily in laminae I-II_o_, less densely in laminae III to dorsal lamina V and only very sparsely in mid-to-ventral lamina V, while some A-MNs innervating glabrous skin provide prominent innervation of the middle and inner parts of lamina II.

Little is known about the function of A-MNs with respect to affective pain and nocifensive behavior. While direct evidence have been lacking, it has generally been thought that one major role is to mediate nociceptive withdrawal reflexes. However, activation of the *Npy2r*^+^ subpopulation of A-MN induced exaggerated withdrawal reflexes, that also involved withdrawal of the contralateral hind paw, and often guarding and jumping; only concurrent activation of low-threshold Aβ-fiber mechanoreceptors (A-LTMRs) would yield an appropriate, well-coordinated reflex^14^. Nevertheless, here we found that a broader activation of A-MNs did not induce an exaggerated reflex but instead a much more restricted and precise reflex of the stimulated paw, with only repeated stimulation evoking licking or guarding. Although we cannot exclude that a small population of A-LTMRs may have been targeted in our mice which could influence these observations, we found neither any innervation indicating association with specialized receptor structures associated with A-LTMRs in the glabrous skin, such as Meissner corpuscles or Merkel cell complexes, nor responses of targeted DRG neurons to low-threshold mechanical stimulation in *in vivo* Ca^2+^ imaging. Thus, it is clear that activating A-MNs in glabrous skin is sufficient to induce well-defined and precise nociceptive withdrawal reflexes.

C-nociceptors, which have also been widely implicated in nociceptive reflexes^15, 16, 18, 40^, were not able to rescue mechanical NWRs in a human Aβ-deafferented individual. Similarly, thresholds for mechanically evoked NWRs were strongly increased in mice where A-MN signaling had been abrogated. In a previous study, optogenetic activation of MrgprD^+^ C fibers evoked paw withdrawal with a latency exceeding 700 ms, while activation of TRPV1-lineage fibers (which include C- and but also some A-nociceptors) yielded withdrawal latencies of ∼150 ms^41^. This is substantially slower than the latency of ∼50 ms observed for pin-prick evoked NWRs^14, 41^. By contrast, the ∼20-ms latency for paw withdrawal induced by optogenetic activation of A-MNs observed here (and previously for *Npy2r*^+^ fibers^14^) appears well-suited to mediate mechanical NWRs, allowing for both nociceptor signaling, spinal processing, efferent signaling and muscle activation within the time course for such reflexes. Thus, we conclude that A-MNs are necessary and sufficient for mechanical NWRs, whereas behavioral responses observed after C-mechanonociceptor activation are not attributed to spinal reflexes but instead depend on supraspinal, potentially including cortical, processes.

In addition to a fundamental role in NWRs, we found that activation of NFH^+^/Na_V_1.8^+^ fibers was also sufficient to trigger affective pain and aversive behavior, in line with what has been reported for a hair follicle-innervating A-MN subpopulation^12^. However, in a parallel study we have recently identified a new A-MN subpopulation characterized by its expression of TrkC, which mediates NWRs but does not appear to be involved in affective pain behavior^42^. Thus, there appears to be a functional division of different cutaneous A-MN populations. Further efforts are needed to untangle the contributions of different subpopulations of A-nociceptor to different aspects of nociception and pain.

An unexpected finding was that prolonged stimulation of NFH^+^/Na_V_1.8^+^ fibers alone induced a marked punctate and dynamic mechanical allodynia. This phenomenon appears to rely on central sensitization, as evidenced by the concomitant prominent phosphorylation of ERK in the spinal cord^43^, but potentially also on an increase in excitability of NFH^+^/Na_V_1.8^+^ fibers themselves, as suggested by the decrease in the light intensity required to activate these fibers by optogenetic means to evoke a withdrawal reflex.

In conclusion, we established intersectional genetic targeting of mouse NFH^+^/NaV1.8^+^ neurons as a powerful tool to study A-MN function and morphology. We showed a fundamental and sufficient role for A-MNs as a broad population in the generation of precise and rapid mechanical nociceptive withdrawal reflexes but also in affective-motivational aspects of pain in mice and humans. We further provided detailed morphological views of A-MN processes, confirming epidermal free nerve endings as the major form of cutaneous innervation and showing unexpectedly restricted central termination in spinal lamina V. Finally, we discovered a novel form of sensitization and allodynia induced by selective stimulation of A-MNs, suggesting an expansive role for these fibers in not only acute nociception but also in long-lasting and pathological pain.

## Acknowledgments

This study was funded by Knut and Alice Wallenberg Foundation, project no. 2019.0047 (ML, MCL, WSJ, and HO), a Knut and Alice Wallenberg Foundation Fellowship (MS), Swedish Research Council (MS, SSN), and the Swedish Brain Foundation (ML, SSN). We thank Dr. Joost Wiskerke at the Technical Platform for Optogenetics and Optical Imaging at the Center for Systems Neurobiology, Linköping University and Drs. Maria Ntzouni and Vesa Loitto at the Microscopy Core Facility at the Medical Faculty, Linköping University for technical assistance and use of equipment.

## Author contributions

ML conceived the study. JCYC, OLM, FMF, AA, OT, SSN, MCL, MS and ML designed the experiments. JCYC, OLM, FMF, AA, LK, KH and ML performed animal experiments and analyzed data from these. OT, JC, and SSN collected and analyzed data from the human subjects. HO, WSJ, SSN, MCL, MS and ML provided supervision. ML wrote the manuscript with feedback from all authors.

## Methods

### Animals

The previously reported NFH^CreERT^^2^ ^23^ and Na_V_1.8^FlpO^ ^24^ mouse lines were crossed with each other and with dual Cre/Flp-dependent reporter mice to generate intersectional triple-crosses; alternatively, NFH^CreERT^^2^;Na_V_1.8^FlpO^ mice were used for adeno-associated virus (AAV) injections (see below). The following reporter mice were used: R26-LSL-FSF-ReaChR-mCitrine (JAX #024846; here called ReaChR mice), ROSA26DR-Matrix-dAPEX2 (JAX #032764; here called APEX2 mice)^44^, Ai195 (JAX #034112; here called GCaMP7s), RC::FPDi (JAX #029040; here called hM4D_i_)^45^. All reporter mice were obtained from Jackson Laboratory. All animal experiments were approved by the Animal Ethics Committee at Linköping University (permits no. 2439-2021 and 214-2021) or Uppsala tingsrätt (5.8.18-01428 and 5.8.18-09954) and performed in accordance with the EU Directive 2010/63/EU.

### Tamoxifen injection

For NFH;Na_V_1.8;ReaChR, NFH;Na_V_1.8;APEX2, NFH;Na_V_1.8;GCaMP7s, NFH;Na_V_1.8;hM4D_i_, or AAV-injected NFH;NaV1.8 mice, tamoxifen (T5648, Sigma-Aldrich; 15495719, Fisher Scientific) dissolved in corn oil at a concentration of 20 mg/mL was administered at 8–10 weeks of age by either a single injection or by two injections (100 μL, i.p.). The mice were used for further experiments no earlier than 2 weeks after injection.

### Tissue preparation

Adult mice of either sex were anesthetized with i.p. injections of either sodium pentobarbital (100 mg/kg) or a mixture of ketamine (120 mg/kg) and dexmedetomidine (0.5 mg/kg), after which they were subjected to transcardial perfusion using 5 mL phosphate buffer (PB, 0.1 M pH 7.4) followed by 50 mL of 4 % paraformaldehyde (for light microscopy) or 2 % paraformaldehyde and 2.5 % glutaraldehyde (for electron microscopy). Tissue of interest was harvested, post-fixed overnight at 4°C and then stored in 1/10 fixative or PB until further processing.

### Immunofluorescence

Tissue specimens were cryoprotected in 30 % sucrose, embedded in OCT, cut at 15 μm thickness in a cryostat and placed on glass slides. Alternatively, specimens were embedded in 4 % low-melting agarose (Fisher Scientific #10377033) and cut on a vibrating microtome (Campden Instruments 7000smz-2) at 50 μm thickness. Sections were incubated in phosphate-buffered saline (PBS) with 3 % normal goat serum, 0. 5 % bovine serum albumin and 0.5 % Triton X-100 (blocking solution), and then in blocking solution with primary antibodies (see Supplemental Table 1) overnight at room temperature. Sections were next rinsed and incubated in solution containing appropriate fluorophore conjugated secondary antibodies (all from Life Technologies, at 1:500 dilution). For detection of isolectin B4 (IB4) binding sites, biotinylated IB4 (Life Technologies, I21214) was added to the primary antibody solution at 1:1,000 dilution, and streptavidin-Alexa Fluor 488 (Life Technologies, S11223) added to the secondary antibody solution at 1:500 dilution. In some cases, cell nuclei were stained using DAPI or SYTOX Deep Red (ThermoFisher Scientific, S11380). Sections were coverslipped with Prolong Diamond, Prolong Glass or SlowFade Diamond (Life Technologies) and imaged using Zeiss LSM800 (10x/0.3, 20x/0.8, 40x/1.3 and 63x/1.4 objectives) or Leica Stellaris 5 (25x/0.95 and 63x/1.4 objectives).

### In situ hybridization

In situ hybridization was based on the RNAScope protocol for frozen fixed tissue sections. DRG cryostat sections (15 μm thickness) were first dehydrated and pretreated with 0.3% hydrogen peroxide and protease IV before the incubation in *Scn10a* probe solution. After probe incubation, the sections were washed using probe wash buffer. For amplification, the sections were then incubated in *Amp1*, *Amp2*, *Amp3*, HRP and OPAL 620 (1:1500 in TSA buffer) solutions. The amplification was stopped by incubation in HRP-blocker solution. After coverslipping with Prolong Diamond, the sections were imaged in a Zeiss LSM800 confocal microscope.

### Image analysis

To measure the soma size of DRG neurons, complete z-stacks (24–33 optical sections at 0.53 μm separation, 24 μm pinhole) of each DRG section were obtained using a Zeiss LSM800 confocal microscope and a 20x/0.8 objective, and analyzed using Fiji. The maximal cross-sectional area of each DRG neuron in the z-stack was determined. Neurons where the largest cross-sectional area was observed in the first or last optical section of the z-stack were excluded. To measure the axon size of sciatic nerves, each sciatic nerve section was imaged using a Zeiss LSM800 confocal microscope and a 20x/0.8 objective. Axon cross-sections with a circularity (defined as 4*π* × *area*/*perimeter^2^*) greater than 0.75 were selected, and their areas were analyzed using Fiji.

### Electron microscopy

Lumbar spinal cord from triple heterozygous NFH;Na_V_1.8;APEX2 mice (4 females, 13-16 weeks old) was embedded in 4 % low-melting agarose and sectioned on a vibrating microtome into transverse 150 µm sections. For APEX2 histochemistry, sections were pre-incubated in 3,3’-diaminobenzidine (DAB; Vector Laboratories) without H_2_O_2_ for 30 min, and then incubated in DAB with H_2_O_2_ for 90 min. The tissue was then osmicated in 1 % OsO_4_, counterstained *en bloc* with 1 % uranyl acetate (Electron Microscopy Sciences) in 50 % ethanol, dehydrated in a graded series of ethanol and embedded in Durcupan ACM (Sigma-Aldrich #44610) using a standard protocol. Ultrathin sections (70 nm thickness) were cut and placed on single-slot copper grids. Some sections were counterstained using 2 % uranyl acetate in H_2_O and 0.5 % lead citrate in H_2_O, while others were used with *en bloc* staining only. Sections were examined in a JEOL JEM1400 Flash transmission electron microscope at 80 kV.

### *In vivo* imaging

*In vivo* Ca^2+^ imaging of DRGs were performed in NFH;Na_V_1.8;GCaMP7s mice essentially as previously described^23^. Each mouse was anesthetized with isoflurane (4 % induction, 1.5 % maintenance) and transferred to a custom surgical platform equipped with a heating pad to maintain body temperature. The dorsal aspect of the spinal cord was surgically exposed between the T12-L1 vertebrae and stabilized with spinal clamps. L4 DRGs were exposed using a dental drill, after which the mouse was transferred to the stage of a custom light microscope (Thorlabs Cerna) equipped with a 4x/0.28 air objective (Thorlabs). GCaMP7s fluorescence images were acquired for 40 second epochs at 5 Hz with an sCMOS camera (Sona 4.2, Andor) using a standard green fluorescent protein (GFP) filter cube. A battery of stimuli was applied to the glabrous skin of the ipsilateral hind paw. The stimuli included gentle brush (using a puffed-up cotton bud), a set of von Frey filaments (0.16 g, 0.60 g, 1.40 g, 8.00 g, 26.0 g), pinblock, and temperature (22°C → 48°C → 22°C). The pinblock stimulus was applied using a resin 3D printed block with an evenly spaced array of 3 × 10 pins; the pin tips had a diameter of ca 100 µm and were spaced 2 mm apart. The hind paw was placed on a 3D printed flexible cushion that limited total downward force to around 3 N. Analysis of Ca^2+^ imaging was performed as previously described^12^. Regions of interest (ROI) of responding cells were outlined in Fiji and the relative change of GCaMP7s fluorescence (ΔF/F) calculated. Background signal (*e.g*., from out-of-focus tissue and neighboring cells), was removed by subtracting the fluorescence of a donut-shaped area surrounding each ROI using a custom MATLAB script. The response threshold was set to 10 %.

### Ex vivo optogenetics and electrophysiology

#### Conduction velocity measurements

NFH;NaV1.8;ReaChR mice were anesthetized with 0.5 mL isoflurane for 1-2 min and euthanized by cervical dislocation. The vertebral column of the mouse was dissected, and lumbar dorsal root ganglia (DRG) along with the connected dorsal roots were carefully extracted in a continuously oxygenated (95 % oxygen, 5 % carbon dioxide) ice-cold cutting solution, which was composed of (in mM): 93 *N*-methyl-D-glucamine, 2.50 KCl, 1.20 NaH_2_PO_4_, 30 NaHCO_3_, 20 HEPES, 25 glucose, 5 sodium ascorbate, 2 thiourea, 3 sodium pyruvate, 10 MgSO_4_, 0.5 CaCl_2_. The extracted root-connected DRGs were then incubated for 30 minutes at 36°C in continuously oxygenated (95 % oxygen, 5 % carbon dioxide) aCSF(mM): 126 NaCl, 2.5 KCl, 1.25 NaH_2_PO4, 26 NaHCO_3_, 10 glucose, 1.5 CaCl_2_, 1.5 MgCl_2_. Thereafter, the root-connected DRG was placed in a recording chamber, and continuously perfused with the same oxygenated (95 % oxygen, 5 % carbon dioxide) aCSF, where the temperature was kept at 34-36°C using a HPT-2 heated perfusion tube (ALA Scientific Instruments Inc.) controlled by a Scientifica temperature controller (Scientifica, UK). The DRG was electrically stimulated with a suction pipette connected to an A365 Stimulus Isolator (World Precision Instruments), and ReaChR expressing DRG neurons were optogenetically activated by a blue light (415nm) from a fluorescent LED light source (CoolLED system, Andover, United Kingdom) through a 10x water immersion objective (LUMPlan FI, 0.90 numerical aperture (NA), Olympus). The activated compound action potentials (CAP) were then measured at the distal end of the dorsal root via a suction pipette (0.5 MΩ) filled with the same aCSF as above. All signals were amplified with a MultiClamp 700B amplifier (Molecular Devices, San Jose, CA), digitalized at 20 kHz with Digidata 1440A (Molecular Devices), low pass filtered at 10 kHz, acquired in WinWCP software (Dr. J. Dempster, University of Strathclyde, Glasgow, United Kingdom), analyzed with Clampfit 11.2 (Molecular Devices).

#### ReaChR activation kinetics

To determine the activation latency of ReaChR, DRGs were collected using the same procedure as described above. DRGs were then immediately incubated in continuously aerated (95 % oxygen, 5 % carbon dioxide) DMEM (Gibco) cell media containing 0.25 % collagenase type IV (Thermofisher) for 1 hour at 36°C (adapted from ^46^). After the incubation, the DRGs were gently dissociated using a fire polished glass pipette. The cell suspension was then centrifuged at 6000 rpm for 5 minutes and the supernatant was carefully removed. The cells were resuspended in the aCSF solution and transferred onto a glass cover slip in a Petri dish filled with continuously oxygenated aCSF solution. A minimum of 2 hours was given for the cells to attach onto the cover slip. Afterward, the Petri dish was transferred to the microscope (same as above). ReaChR positive cells were identified by endogenous mCitrine fluorescence. Cell-attached current-clamp recording was performed using a recording pipette (7-9 MΩ) filled with the same aCSF solution as above. Electrophysiological recordings were acquired as above.

### Behavioral assays

#### Optogenetically evoked paw reflex latencies

Mice were placed individually in an observation cubicle (W×D×H 9×5×5 cm^3^) made of transparent acrylic glass with a 5 mm thick floor, and stimulated on the plantar hind paw. The mice were video recorded from a lateral view using a high-speed camera (a customized C-mount Sony RX0 II (Backbone, Kanata, Canada)) at 1000 frames/s (yielding a temporal resolution of 2 ms). Optogenetic stimulation was applied to the hind paw (460 nm, single 5 ms pulse) via the tip of a 960 µm/NA 0.63 optical fiber patch cord manually placed immediately below the floor beneath the hind paw.

#### von Frey assays

Mice were placed in a cubicle (W×D×H 9×5×5 cm^3^) with clear acrylic walls and a mesh floor. After 15 min habituation, the mechanical paw withdrawal threshold was assessed using manual von Frey filaments applied to the plantar hindpaw. The mechanical von Frey threshold was determined using the simplified up-down method^47^.

To determine von Frey thresholds during chemogenetic inhibition, triple heterozygous NFH;Na_V_1.8;hM4D_i_ mice and control littermates (bearing the same alleles except being double negative for Na_V_1.8^FlpO^), were on Day 1 placed in the mesh floor cubicle and the baseline threshold determined. On Day 2, the animals were injected with either saline or CNO (1 mg/kg i.p.), and von Frey threshold assessed 30 min after injection. On Day 3, the animals were again tested after being injected with the solution not administered on Day 2; the order of solutions was individually randomized for each mouse. Treatment order and genotype were unknown to the experimenter at the time of testing.

Mechanical paw withdrawal thresholds were also tested in mice where synaptic transmission from NFH^+^/Na_V_1.8^+^ fibers had been abrogated using TeTxLC. Here, 9 NFH^CreERT2^;/Na_V_1.8^FlpO^ pups (P0-P4) of either sex were briefly anesthetized with isoflurane and injected i.p. with AAV9.Con/Fon-TeTxLC (1 µL of 8.2×10^12^ vg/mL solution diluted 1:10 in saline, for a final injected volume of 10 µL). Each pup received two injections, either at P0 and P3 or P2 and P4. The TeTxLC plasmid was constructed by Dr Hendrik Wildner, University of Zürich, and the AAVs produced by the University of Zürich Viral Vector Facility. As controls, 6 littermate mice with the same genotype were instead injected with an unrelated AAV (AAV9.Con/Fon-syp-mCherry, from the same source; 1 µL of 1.1×10^13^ vg/mL diluted 1:10 in saline) at P0 and P3. After tamoxifen injections at 6-10 weeks age, the mice were subjected to von Frey assays as above.

#### Hot plate assay

On two separate days, NFH;Na_V_1.8;hM4D_i_ mice and control littermates were injected with saline or CNO (1 mg/kg i.p.). After 30 min, the mouse was placed on a hot plate (IITC Life Science analgesia meter) set at a constant temperature of 50°C, and the latency to jumping or paw licking/flicking measured. The mouse was immediately removed from the plate after a nocifensive reaction was observed. Each mouse was subjected to three trials (minimum inter-trial interval 15 min) and the average latency calculated. On the second day, the mouse was administered the solution that it did not receive the first day. The order of saline and CNO injections was individually randomized. The experimenter was blind to both treatment order and genotype.

#### Real-time place preference

On Day 1 (pre-test), triple heterozygous NFH;Na_V_1.8;ReaChR mice and littermates (same genotype except being FlpO negative) were placed in a two-chamber clear acrylic glass apparatus (175 × 90 mm^2^ each) connected via a 50 mm wide opening. The floor consisted of 5 mm thick clear acrylic. Mice were video recorded from above using a webcam (Logitech C920) and AnyMAZE software. During the pre-test, the mice were allowed to explore the apparatus freely for 15 min. The chamber in which each mouse spent most of its time during pre-test was defined as its preferred one. On Day 2 (test phase), the mice were again placed in the apparatus for 15 min. Now, one hind paw was continuously tracked using an optic fiber tip (NA 0.63, 960 µm core, attached to a Prizmatix 460 nm LED) manually placed subjacent to the floor, immediately below the hind paw. When the mouse entered the chamber initially preferred during the pre-test, optogenetic stimulation (20 Hz, 5 ms pulses) was switched on automatically via the AnyMAZE software, and remained on until the mouse exited to the other chamber. The preference score *P* was calculated as *P* = 100 × (*t*_*test*_ − *t*_*pre*_)⁄*t*_*pre*_, where *t*_*test*_ is the time spent in the light-on chamber during the test phase and *t*_*pre*_ the time spent in the same chamber during pre-test. The experimenter was nominally blind to the genotype of the mice throughout the experiment; however, because triple heterozygous mice but not FlpO-negative littermate mice showed overt reflexive and escape behavior in response to optogenetic stimulation, complete blinding was not possible.

#### Facial expression and nocifensive behavior assays

Triple heterozygous NFH;Na_V_1.8;ReaChR mice and littermates were placed in a transparent acrylic glass cubicle (9 × 5 × 5 cm^3^). The lateral view of the cubicle was recorded using a color FLIR camera (Blackfly BFS-U3-23S3C-C) with a 1:1.8/4 mm Basler lens (C125-0418-5M) at 60 frames/second. After 10 min of habituation, baseline was recorded for 3 min, after which optogenetic stimulation was applied to the plantar hind paw through the transparent floor via a fiberoptic ferrule (0.63 NA, 960 µm; 5 ms pulses at 20 Hz using a Prizmatix 460 nm LED or a Doric Lenses 595 nm LED). After stimulation, a 3 min recovery period was also recorded. Facial expression analysis was performed as described^32^. For analysis of nocifensive behavior, the first 60 s of the optogenetic stimulation period was analyzed using Noldus The Observer XT 15 software with respect to paw withdrawal, jumping (on hind limbs or all four limbs, counted separately), paw licking and paw guarding. Event counts were obtained for each behavior, as were the durations of licking and guarding behaviors.

#### Prolonged optogenetic stimulation

NFH;Na_V_1.8;ReaChR mice of either sex were, during isoflurane anesthesia, subjected to optogenetic stimulation (465 nm, 20 Hz, 5 ms pulses) for 5 min to the left plantar hind paw. After 10 min of recovery from anesthesia in a cubicle with mesh floor, a soft brushing stimulus (using a puffed-up cotton bud) was applied to the hind paw; this was repeated three times and the response frequency recorded. Next, the mechanical withdrawal threshold was tested using von Frey filaments (as above) in ipsi- and contralateral hind paws 10 min after recovery from anesthesia. Finally, the light intensity needed to evoke a paw withdrawal was tested. To assess this threshold, 5 ms pulses (20 Hz, NA 0.63, 960 µm core, attached to a 595 nm LED from Doric Lenses) were applied to the hindpaw from below the mesh floor at five set intensity levels, from the lowest intensity to the first that initiated a response. Because the measured power from the LED changed somewhat between experimental sessions, absolute threshold values varied slightly between mice.

A separate group of mice were used for immunofluorescence of extracellular signal-related kinase 1/2 phosphorylated at Thr^202^/Tyr^204^ (pERK), a marker of central sensitization in the spinal cord^43^. Here, immediately after the prolonged optogenetic stimulation and under continued isoflurane anesthesia, the mice were subjected to perfusion fixation using 4 % paraformaldehyde, after which spinal cord tissue was processed for immunofluorescence as above.

### Human reflex electromyography and psychophysics

#### Participants

28 healthy controls (HCs), 18 to 40 years old (females, 18, males, 10), were originally recruited from an existing database at the Center for Social and Affective Neuroscience (Linköping University) and advertisements on social media sites. Exclusion criteria were diabetes, muscular, skeletal, skin, or neurological diseases, analgesic and/or psychoactive medication. Due to insufficient data (e.g, missing NWR thresholds or incomplete data collection), a total of 22 controls were included in the analysis. The individual with Aβ deafferentation (male, 70 years) is well-characterized and suffers from a rare sensory ganglionopathy syndrome^33^. Because this individual belongs to an older age group, an age-matched individual (female, 77 years) was used for comparison in NWR and thermal threshold testing. Written informed consent was obtained from all individuals before the start of the experiment and all data processing followed the GDPR guidelines. The study was approved by the ethics committee of Linköping University (dnr 2020-04207) and Liverpool John Moores University (14/NSP/039).

#### Mechanical reflex elicitation

To evoke the reflex mechanically, a custom-built device delivered a single pinprick. A solenoid motor (Mecalectro 819AB83) launched a pin (1 mm diameter) to a maximum travel distance of 9 mm before retracting after each stimulus. An optical sensor placed next to the pin registered the time the pin passed through its hole to signal stimulus onset. The stimulus had a bandwidth up to ∼20 Hz, amplitudes up to ∼20 µV, and a pulse width of ∼400 µs. The mechanical indentation was adjusted by changing the voltage of the power supply (EA-PS 2342-10 B), which ranged from 5 to 35 V. The power sequence was controlled using EasyPS2000 (version 2.05, with LabView runtime, 236 MB, Elektro-Automatik, Viersen, Germany). The timing of the stimulus was manually randomized between 3 s and 1 min to avoid habituation of the reflex and maintain the novelty of the stimulus. A silent computer mouse was used to trigger the device, and a physical partition blocked visual cues.

#### Experimental setup

Participants were seated in a comfortable chair with the right foot hanging freely at a 90-130° angle. The foot was placed on the device platform, at a slight upward angle over a small rectangular hole, allowing the pin to deliver the stimulus just below the anterior lateral eminence of the foot sole, an area corresponding to the reflex receptive field of the tibialis anterior muscle^48^. Two adhesive electrodes were placed 2 cm apart on the scratched and degreased skin over the belly of the tibialis anterior muscle on the leg, as anode and cathode, and a third electrode was placed over the kneecap as reference (Kendall ECG electrodes 57×34mm, Medtronics, USA).

The foot soles were examined after each stimulus to mark the spot where a reflex could be evoked and to ensure that the skin was not sensitized, the absence of redness and any lingering pain or discomfort was confirmed following the stimulus. Reflexes were recorded using PowerLab (16/35 AD Instruments, Oxford, UK), and stored in LabChart (v8.1.16 AD Instruments, Oxford, UK) with a low pass filter of 1 khz and a high pass filter of 0.3 Hz, a range of 1 mV, sampled at a rate of 20 000 Hz with an acquisition delay of 160 µs. A clear deviation from baseline, evoking a reflex response with a z-score >1 was included in the analysis. Reflex latencies were further calculated, post-acquisition, in MatLab (R2021b, MathWorks Inc, Natick, Massachusetts). A time analysis window of 40-200ms was used to include both short- and long-latency NWR responses.

#### Pain and reflex thresholds

To assess pain intensity, a visual analog scale (VAS) was used, ranging from 0 representing “no pain” (to the far left) to 10 representing “worst imaginable pain” (to the far right). The mechanical indentation was increased stepwise, in 1V increments, until the participant reported a painful sensation. This was taken as the pain threshold, i.e. defined as the voltage at which the participant first rated something above 0 on the scale. Pain ratings were further analyzed, post-acquisition, in MatLab (R2021b, MathWorks Inc, Natick, Massachusetts). Reflex thresholds were defined as the stimulus voltage at which the NWR was elicited three times at the same location on the foot sole. All stimulations were delivered during a relaxed EMG state, without any visible muscle contractions. Once thresholds were established during baseline, stimulus intensities at pain and NWR thresholds were applied a minimum of 3 times during block and at least once during recovery.

#### Temperature detection thresholds

As Aδ and C fibers mediate cold and warm sensations, respectively, in humans^49, 50^, quantitative thermal testing (TCSII, Strasbourg, France) was performed to determine cold and warm detection thresholds (CDT and WDT) at a rate of 1°C/s in a series of 4 trials. All baseline temperatures were set to the individual’s skin temperature and CDT and WDT thresholds are reported as baseline temperature – threshold temperature (or delta thresholds ΔCDT and ΔWDT).

#### Nerve conduction velocity testing in the Aβ-deafferented individual

To ensure that any effect on the NWR was not due to slow conducting efferent fibers, nerve conduction velocity testing was conducted on the individual with Aβ deafferentation using standard laboratory equipment (Nicolet EDX, Natus Neurology Incorporated, Middletown Wisconsin, USA). Recording electrodes (Nicolet Biomedical, EMG surface electrode, Cephalon, Denmark) were attached to the belly of the tibialis anterior, extensor digitorum brevis, and the abductor digiti minimi muscles 2 cm apart and connected to a stimulator (Nicolet AT2+6 amplifier, Natus Neurology Incorporated, Middletown Wisconsin, USA). Stimulating electrodes (Nicolet Biomedical, EMG surface electrode, Cephalon, Denmark) were placed over the ulnar, peroneal, and tibial nerve. Conduction velocities (CV) were automatically calculated between a distal and proximal site on all nerves by the computer software (Synergy CareFusion EDX, V.20.0, 2010, Middleton Wisconsin, USA).

### Statistics

All descriptive statistics and statistical tests were performed in GraphPad Prism. Normal distribution of data was assessed using the Shapiro–Wilk normality test. Measures are given as mean ± SD or as median ± interquartiles and range, as appropriate.

---

**Supplemental Table 1.**
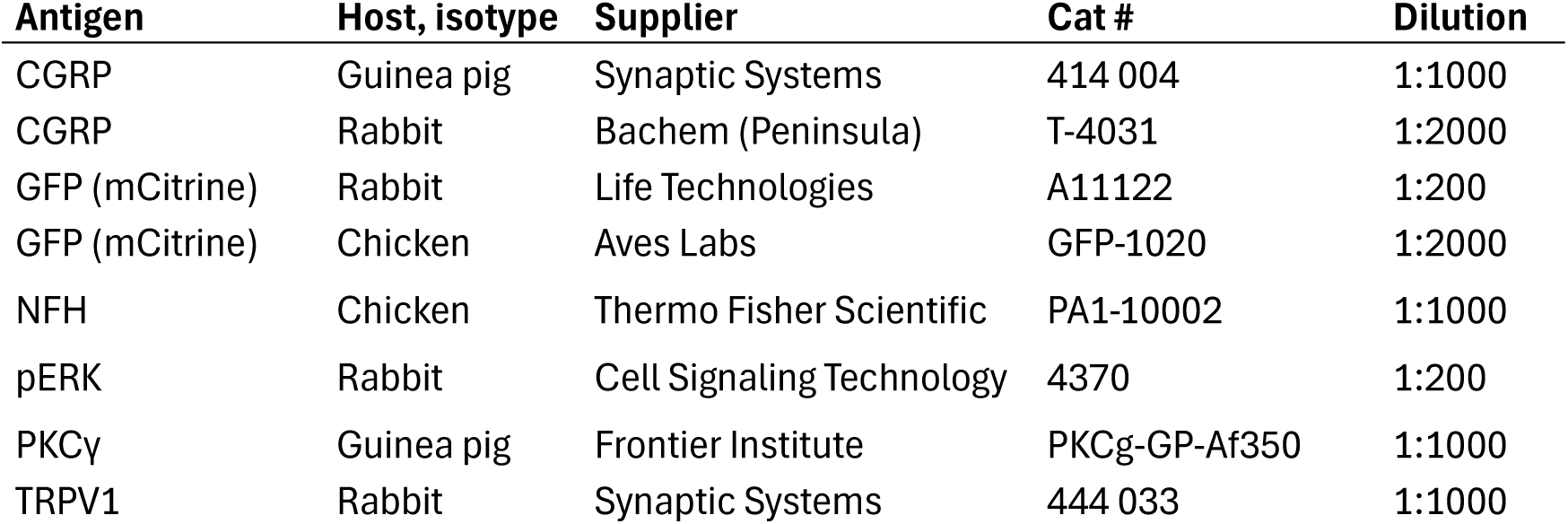
Primary antibodies.

**Figure S1. Measurement of ReaChR activation kinetics in DRG neurons. (a)** Schematic of the setup. Primary mCitrine^+^ DRG neurons isolated from NFH;Na_V_1.8;ReaChR mice were subjected to optogenetic stimulation while patch-clamped. (b), an example voltage trace shows a response (arrow) to a 2 ms light pulse (blue line). The latency was calculated from the start of the light pulse.

**Figure S2. Assessment of peripheral nerve fiber function in the Aβ-deafferented individual. (a)** Temperature detection thresholds. Cold (**CDT**) or warm (**WDT**) temperature detection thresholds did not differ between the Aβ-deafferented individual and the age-matched individual, suggesting that the absence of an NWR in the Aβ-deafferented individual was likely not due to non-functioning Aδ or C fibers. ns, *p* = 0.49 (CDT) and *p* = 0.08 (WDT); unpaired t-test with Welch’s correction. **(b)** Nerve conduction velocity testing. Amplitudes in the upper limb were normal and conduction velocities (CV) were within normal range. Amplitudes were lower in the legs, with CV slightly under normal values. This could have possibly led to slower NWR latencies, but it seems unlikely that mild motor CV impairment leads to a complete NWR abolishment. **ADM**, abductor digiti minimi.

