## Supplementary figures and images for "Central role for fast nociceptors in mechanical nocifensive behavior and sensitization"

### Fig. S1

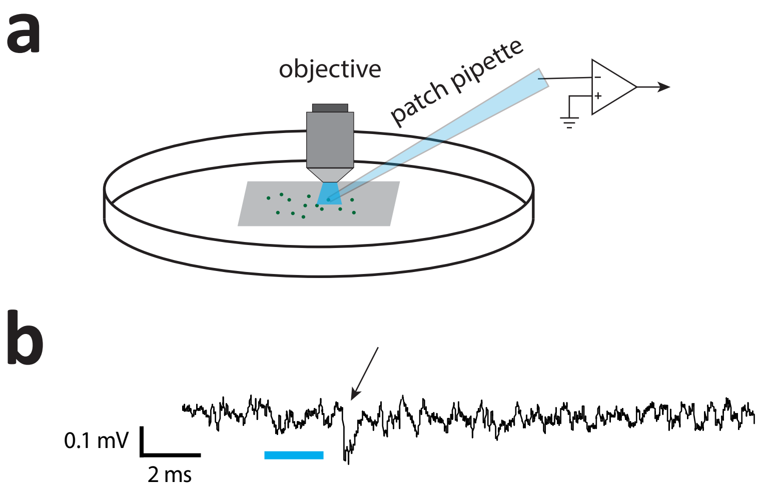


**Figure S1**
