## Supplementary material for "Central role for fast nociceptors in mechanical nocifensive behavior and sensitization": Fig. S2

a

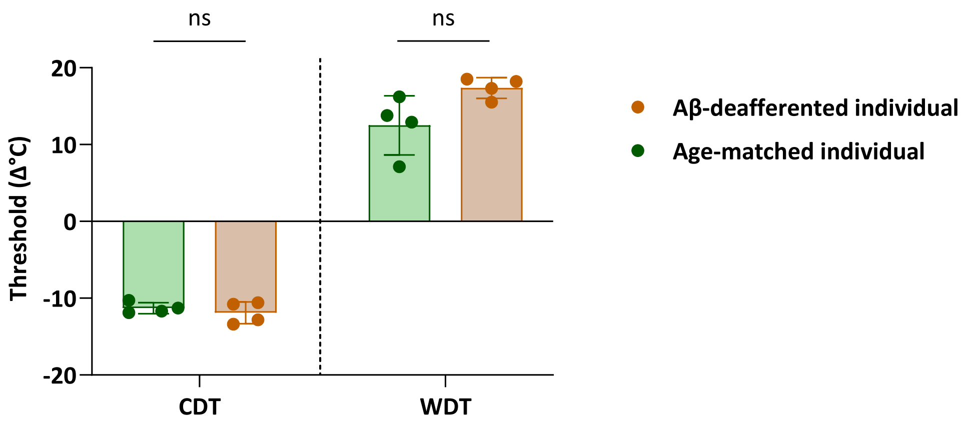


b

| | Latency (ms) | Amplitude (mV) | Amplitude (%) | Conduction velocity (m/s) |
|---|---|---|---|---|
| **R Ulnar – ADM fractioned** | | | | |
| Wrist | 3.28 | 9.4 | 100 | |
| Under Elbow | 7.86 | 4.1 | 43.8 | 44.7 |
| Above Elbow | 10.42 | 7.8 | 82.9 | 37.2 |
| Axil | 13.91 | 5.8 | 61.5 | 45.9 |
| **R Peroneal** | | | | |
| Ankle | 6.56 | 2.8 | 100 | |
| Knee | 18.13 | 2.3 | 82.9 | 34.6 |
| **R Tibial** | | | | |
| Ankle | 6.46 | 1.3 | 100 | |
| Knee | 18.65 | 0.8 | 59 | 36.1 |

**Figure S2**
